# Yeast Models Of Phosphomannomutase 2 Deficiency, A Congenital Disorder Of Glycosylation

**DOI:** 10.1101/414862

**Authors:** Jessica P. Lao, Nina DiPrimio, Madeleine Prangley, Feba S. Sam, Joshua D. Mast, Ethan O. Perlstein

## Abstract

Phosphomannomutase 2 Deficiency (PMM2-CDG) is the most common monogenic congenital disorder of glycosylation (CDG) affecting at least 800 patients globally. PMM2 orthologs are present in model organisms, including the budding yeast *Saccharomyces cerevisiae* gene SEC53. Here we describe conserved genotype-phenotype relationships across yeast and human patients between five PMM2 loss-of-function missense mutations and their orthologous SEC53 mutations. These alleles range in severity from folding defective (hypomorph) to dimerization defective (severe hypomorph) to catalytic dead (null). We included the first and second most common missense mutations – R141H, F119L respectively– and the most common compound heterozygote genotype – PMM2^R141H/F119L^ – observed in PMM2-CDG patients. Each mutation described is expressed in haploid as well as homozygous and heterozygous diploid yeast cells at varying protein expression levels as either SEC53 protein variants or PMM2 protein variants. We developed a 384-well-plate, growth-based assay for use in a screen of the 2,560-compound Microsource Spectrum library of approved drugs, experimental drugs, tool compounds and natural products. We identified three compounds that suppress growth defects of SEC53 variants, F126L and V238M, based on the biochemical defect of the allele, protein abundance or ploidy. The rare PMM2 E139K protein variant is fully functional in yeast cells, suggesting that its pathogenicity in humans is due to the underlying DNA mutation that results in skipping of exon 5 and a nonfunctional truncated protein. Together, these results demonstrate that yeast models can be used to characterize known and novel PMM2 patient alleles in quantitative growth and enzymatic activity assays, and used as patient avatars for PMM2-CDG drug screens yielding compounds that could be rapidly cross-validated in zebrafish, rodent and human organoid models.

## Introduction

Phosphomannomutase 2 deficiency (PMM2-CDG) is the most common congenital disorder of glycosylation (Ferreira *et al.*, 2018), and was originally described in the literature as CDG1a/CDG1/Jaeken Syndrome (Matthijs *et al.*, 1997). PMM2-CDG is an autosomal recessive, multi-organ monogenic disease displaying variable clinical progression and presentation. It is caused by an underlying enzymatic deficiency in PMM2 leading to cell-autonomous defects in the production of N-linked glycoproteins. Primarily affected organs include brain, liver, gastrointestinal tract, heart, kidney, and most PMM2-CDG patients have intellectual disability and developmental delay (Schiff *et al.*, 2017). PMM2-CDG is a global disease but the exact worldwide incidence of PMM2-CDG is unknown. There are currently no FDA-approved therapies for PMM2-CDG.

The yeast ortholog of PMM2, SEC53, was originally recovered as a *sec* (short for secretory) complementation group in forward genetics screens in the early 1980s. These studies culminated three decades later in the 2013 Nobel Prize in Medicine and Physiology (Ferro-Novick *et al.*, 1983; Kepes & Schekman, 1988). All PMM2 orthologs including SEC53 isomerize mannose-6-phosphate (M6P) to mannose-1-phosphate (M1P), and are activated by glucose-1,6-bisphosphate (G16) and mannose-1,6-bisphosphate (M16) (Pirard *et al.*, 1999). M1P is a precursor of GDP-mannose, which in turn is required for the production of dolichol phosphate mannose and then lipid-linked oligosaccharides (LLO) in the lumen of the endoplasmic reticulum. LLOs are substrate by a cascade of enzymes that are individually linked to a CDG (Panneerselvam & Freeze, 1996). The oligosaccharyltransferase (OST) complex executes the final steps of N-linked glycosylation by transferring the glycan from the LLO donor to asparagine residues of nascent proteins (Cherepanova *et al.*, 2016). *sec53* yeast mutants grown at the non-permissive temperature accumulate hypoglycosylated forms of secretory proteins, leading to intracellular buildup of normally secreted proteins and ultimately cell death. Over-expression of human PMM2 in a *sec53* temperature-sensitive mutant rescued lethality (Hansen *et al.*, 1997).

PMM2 activity has been studied *in vitro* using recombinant enzyme expressed in bacteria and in patient-cell-derived assays. PMM2 protein variants were expressed in *E. coli* and displayed enzymatic activities between 0% and 100% of control levels, with most missense alleles exhibiting activity between 16-54% of control levels (Kjaergaard *et al*., 1999; Vega *et al.*, 2011). The limitation of this model system is that it fails to replicate heterodimer formation, which is almost always the case in PMM2-CDG patients. Patient fibroblasts, which are fast-dividing cells, have residual activity between 35%-70% of control levels, and higher passage number is associated with increased basal PMM2 activity (Grunewald *et al.*, 2001). Patient-derived leukocytes, which are terminally differentiated cells, have less background contaminating PMM2 enzymatic activity and show clear separation between cases and controls (Grunewald *et al.*, 2001). Similarly, induced pluripotent stem cell (iPSC) clones have been derived from patient fibroblasts and they also display proteome-wide hypoglycosylation (Thiesler *et al.*, 2016).

Attempts to create a viable PMM2 mouse model have been stymied at multiple turns. PMM2 knockout mice are early embryonic lethal (Thiel *et al.*, 2006). Several years ago, a PMM2^R141H/F119L^ knock-in mouse model was generated, and they exhibit heterozygous mice have variable lethality starting prenatally with few postnatal escapers (Chan *et al.*, 2016). Challenges in creating viable PMM2 mutant mice have encouraged parallel efforts to create disease models in other model organisms such as nematodes, flies and zebrafish. The nematode PMM2 ortholog (F52B11.2) is uncharacterized, and the first *Drosophila* model of PMM2 deficiency were published two years ago (Parkinson *et al.*, 2016). Flies homozygous for strong loss-of-functions alleles for *pmm2* die during the larval stage. Flies in which the expression of *pmm2* is “knocked-down” using RNA interference in neurons have both morphological and physiological synaptic defects at the neuromuscular junction, and locomotor defects. A morpholino-induced PMM2 knockdown zebrafish model results in reduced N-linked glycosylation and LLO levels, along with craniofacial and motor defects reminiscent of PMM2-CDG patients (Cline *et al.*, 2012).

The relationship between PMM2-CDG clinical progression and presentation and PMM2 enzymatic activity measured using recombinant protein variants or in patient cell lysates has been opaque. On the one hand, the absence of R141H homozygotes indicates that similar to other monogenic inborn errors of metabolism, complete PMM2 deficiency is incompatible with human life. This is consistent with the lethality observed in PMM2 deficiency across model organisms (Matthijs *et al.*, 1998; Schollen *et al.*, 2000). R141H is a catalytic dead variant functionally equivalent to a knockout or genetic null. On the other hand, the over-abundance of missense mutations in PMM2 in large exome databases of control populations suggests that there could be heterozygote advantage to 50% PMM2 enzyme levels (Citro *et al.*, 2018). That would also explain the high incidence of novel missense mutations *in trans* with R141H.

Hypoglycosylation and varying degrees of lethality are common phenotypes observed across all PMM2 animal models. These results suggest that the minimum threshold of PMM2 enzyme activity required for viability may vary depending on the physiological state of the cell, cell differentiation status and stage of organismal development. The differential susceptibility of cell types may be attributed to proteome-wide hypoglycosylation and/or loss of glycosylation of specific regulatory or housekeeping glycoproteins or a combination of both. The full extent of PMM2-CDG pathophysiology has not been elucidated, perhaps in part because model organism efforts have not been coordinated and focused on specific patient mutations. We turned to budding yeast to create foundational PMM2-CDG patient avatars for multiple uses, starting with growth-based phenotypic drug screens to rapidly identify therapeutic options tailored to specific patient mutation combinations.

## Methods

### Strains and plasmids

All strains used in this study are in S288c background (Supplemental Table 1). Strains were grown in SC or SC drop out media (Sunrise) + 2% dextrose at 30°C unless otherwise noted. Standard genetic procedures of transformation and tetrad analysis were followed to construct strains. SEC53 rescue plasmid was generated by cloning the SEC53 promoter, open reading frame, and terminator sequences into the episomal pRS316 containing the URA3 selectable marker. SEC53 variants were generated by cloning the relevant promoter, GFP or SEC53 gene, and CYC1 terminator sequences into an integrating plasmid containing the LEU2 selectable marker and integrated at the SwaI restriction site into the HO locus. For PMM2 variants, the SEC53 ORF was replaced with a codon-optimized PMM2. Plasmids were generated by Next Interactions or GenScript. Strains were generated by Next Interactions or in-house.

### Growth assay

Cells were picked from plates and resuspended in SC media to OD_600_ = 1.0, then serial diluted into 50 µL SC+FOA media in 384-well plates at 10^-1^, 10^-2^, and 10^-3^. Plates were incubated at 30°C and absorbance reading at 600 nM were measured by a plate reader (Molecular Devices SpectraMax M3) at the indicated time points. Plates were vortexed briefly to resuspend cells prior to plate readings. 5-floroortic acid was purchased from US Biological and used at a concentration of 1 mg/L.

### PMM2 enzymatic assay

Yeast lysates were prepared from 50 OD_600_’s worth of cells. Cells are washed once in 25 mM KPO_4_ pH 8.0 and resuspended in 600 µL lysis buffer (25 mM Tris-HCl pH 7.5, 1 mM EDTA, 100 mM NaCl, 10 mM β-mercaptoethanol, and 1 Roche protease inhibitor tablet) and an equivalent amount of glass beads. The bead/lysate mixture was vortexed for 7-10 cycles of 2 min vortex and 1 min cooling on ice. Lysis was checked by microscopy to confirm that 80-95% of cells are lysed. The lysates were clarified by 15 mins centrifuge at max speed at 4°C and the supernatant moved to a new tube. For long term storage at -80°C, glycerol was added to 20% of the total volume. Protein concentration was determined with the Qubit protein assay kit (Thermo Fisher Scientific).

PMM2 enzymatic activities were assayed spectrophotometrically at 340 nm by the reduction of NADP+ to NADPH in a reaction mixture incubated at 30°C for 60 mins in a 96-well plate. Cell lysates were added to 200 µL volume in the following reaction: 50 mM HEPES, pH 7.1, 5 mM MgCl_2,_ 0.5 mM NADP+, 10 µg/mL yeast glucose-6-phosphate dehydrogenase, 20 µM glucose 1,6-bisphosphate, 200 µM mannose 1-phosphate, 10 µg/mL phosphoglucose isomerase, and 5 µg/mL phosphomannose isomerase. Glucose 1,6-bisphosphate was used as an activator of PMM2 activity. NADPH formation was calculated from absorbance at 340 nm using the Beer’s Law (absorbance = εLc).

### Drug Repurposing Screen

125 nL of compounds or DMSO were dispensed into 384-well plates using the Echo acoustic dispenser (LabCyte) to achieve a final concentration of 25 µM. 50 µL of 10^-2^ dilution of an OD_600_ = 1.0 or 0.5 yeast cell suspensions were dispensed into the 384-well plates containing compounds or DMSO with a MultiFlo automated dispenser (Biotek). Plates were covered and incubated at 30°C for 16 – 21 hours until OD_600_ reaches ∼0.8. Plates were vortexed briefly to resuspend cells prior to reading.

### Validation

Microsource Spectrum compounds were reordered from MicroSource Discovery Systems and resuspended in DMSO. 5 – 125 nL compounds or DMSO were dispensed into 384-well plates and 50 µL of yeast cell suspensions or media were dispensed by multichannel pipettes to achieve the desired concentration. Plates were covered and incubated at 30°C for 16 – 23 hours until OD_600_ reaches ∼0.8. Plates were vortexed briefly to resuspend cells prior to plate readings.

## Results

### Modeling PMM2 patient alleles in yeast SEC53

PMM2 and SEC53 are orthologs that share 55% identity at the amino-acid level (**Figure 1A**). We generated three common PMM2 disease-causing alleles R141H, F119L, and V231M, and two less well studied pathogenic alleles: E93A and E139K (**Figure 1B**). The positions of these residues are indicated in **Figure 1C**. R141 is in the substrate-binding domain of PMM2 (Andreotti *et al.*, 2015). R141H has no detectable enzymatic activity and never occurs in homozygosity in patients despite being the most commonly observed mutation (Kjaergaard *et al*., 1999). F119 is a component of the hydrophobic core within the dimer interface and is the second most commonly observed mutation (Silvaggi *et al.*, 2006). F119L has 25% enzymatic activity *in vitro* and this deficiency is likely due to its diminished ability to dimerize and decreased dimer stability (Kjaergaard *et al.*, 1999; Pirard *et al.*, 1999; Andreotti *et al.*, 2015). V231 is in the interior of the core domain and a mutation in this residue is detrimental to its native protein structure (Silvaggi *et al.*, 2006; Citro *et al.*, 2018). The folding and stability defect of the V231M allele contributes to its reported reduced *in vitro* enzymatic activity of 38.5% (Kjaergaard *et al.,* 1999; Pirard *et al.*, 1999). Existing data on E93A and E139K variants are limited. E93 directly interacts with R116 *in trans* within the PMM2 dimer and a mutation in this residue likely compromises dimerization (Andreotti *et al.,* 2015). E139K is a result of a 415G>A transition mutation in the genomic sequence that interferes with RNA splicing, causing either skipping of exon 5 to form a partially deleted and nonfunctional protein or a full-length E139K mutant protein (Vuillaumier-Barrot *et al.,* 1999).

**Figure 1.**
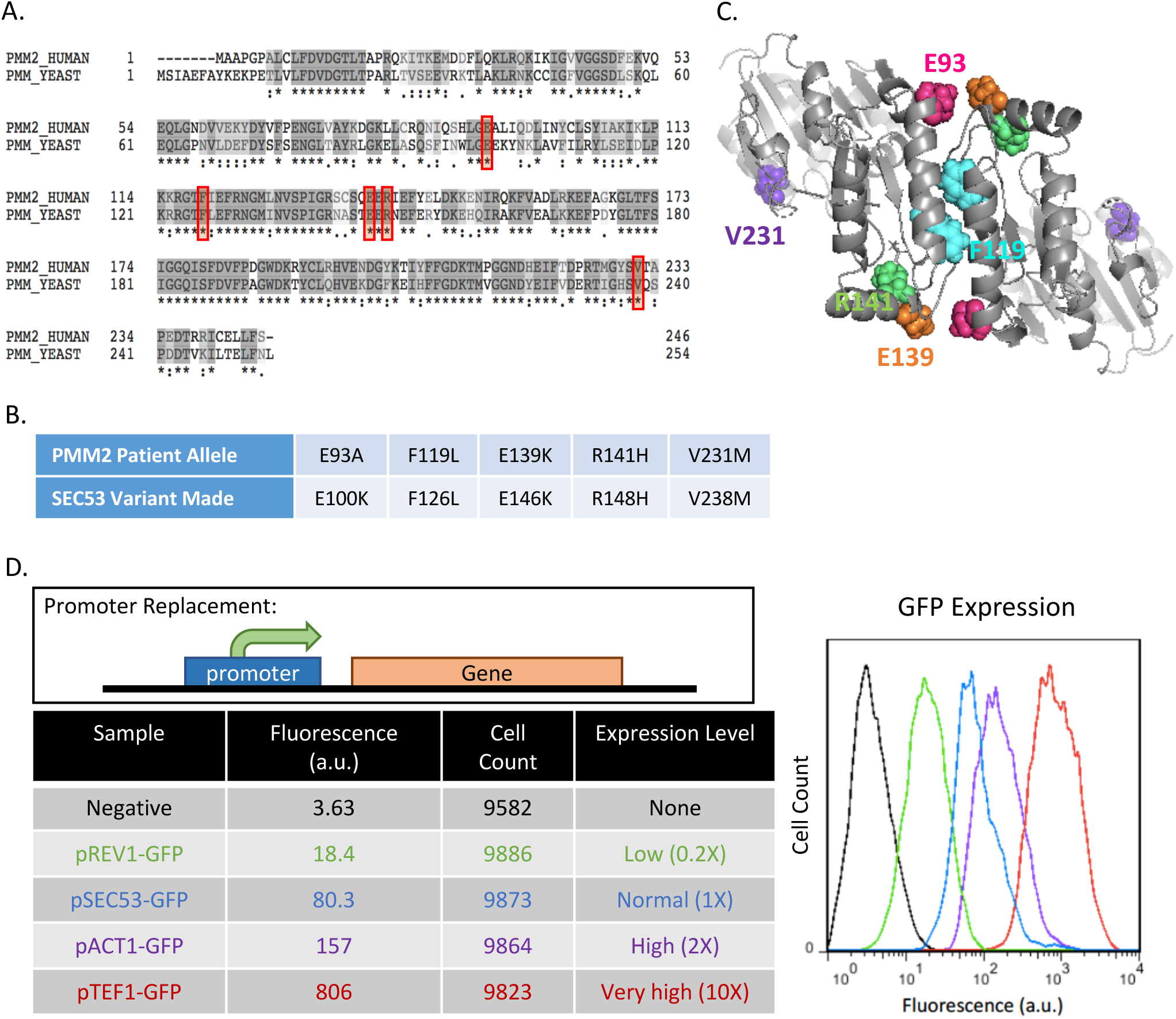
Generating yeast models of PMM2 deficiency. (A) Sequence alignment of phosphomannomutase genes in human (PMM2) and yeast (SEC53). Asterisk (*) indicates an identical amino acid residue and colon (:) indicates similar amino acids. Red boxes show the conserved disease-causing amino acid residues that we’ve modeled. (B) Table showing the PMM2 patient alleles and the equivalent variants we generated in yeast SEC53. (C) Structure of PMM2 dimer highlighting the five modeled residues. Dimer structure was generated from 2AMY in the RCSB protein data bank and courtesy of Dr. Maria Vittoria Cubellis (University of Naples Federico II, Italy). (D) Comparison of promoter strength. Different promoters are used to drive the expression of the gene of interest. GFP is placed under the REV1, SEC53, ACT1, or TEF1 promoter and the fluorescent intensity of GFP is measured by flow cytometry. The graph displays the fluorescence in arbitrary unit (a.u.) against the cell count. The raw numbers are shown in the table along with the expression level relative to the SEC53 promoter.

PMM2 F119L, R139K, R141H, and V231M correspond to SEC53 F126L, E146K, R148H, and V238M, respectively. SEC53 E100K was unintentionally generated, but the residue is conserved in PMM2 and the patient allele is E93A. Some of these variants have low to no detectable enzymatic activity reported in the literature, so we placed the mutants under different promoters to determine if changes in protein abundance affect viability of each variant. The relative strength of the TEF1, ACT1, and REV1 promoters were compared to the native SEC53 promoter by driving expression of the green fluorescent protein (GFP) (**Figure 1C**). Based on fluorescence reading by flow cytometry, we found that relative to the native SEC53 promoter, the strength of TEF1, ACT, and REV1 promoters are 10X, 2X, and 0.2X, respectively.

### Growth of Sec53 variants correlates with enzymatic defects of the variant and promoter strength

To overcome the complication that SEC53 is an essential gene, we placed a wildtype SEC53 copy on a URA3 plasmid that we can conditionally remove by growing cells in 5-fluoroorotic acid (5-FOA). 5-FOA is an analog of uracil that is converted into a toxic intermediate in cells where the uracil biosynthetic pathway is active, which the URA3 marker enables. Each SEC53 variant is then individually integrated at the HO locus of *sec53Δ* cells. The phenotype of each variant is revealed when the wildtype URA3 containing plasmid is counter-selected in media containing 5-FOA.

Increasing the expression level of the hypomorphic alleles improves their growth in every case except the R148H variant and the *sec53Δ* negative control (**Figure 2A, B**). When expressed at native level, the V238M allele is sufficient for growth, but at a slower rate and achieving a lower final yield than wildtype cells (31.8%). Doubling the expression of the V238M allele with the ACT1 promoter restores growth of this mutant: 67% at 20 hours and near wildtype final yield at 24 hours. The F126L allele grows poorly under the endogenous promoter (26.9%), and also grows better when its expression is doubled (56.6%). The relative growth of F126L and V238M is consistent with their reported residual *in vitro* enzymatic activity (Pirard et al., 1999). Over-expression of F126L and V238M alleles under the TEF1 promoter completely rescues these cells. E100K is viable only under the ACT1 (16.2%) and TEF1 (66.5%) promoters, which indicates the severity of this mutation compared to the other variants. On the other hand, the R148H null allele is not compatible with growth at any promoter strength and phenocopies the genetic null *sec53Δ*. Together, these data can be fully explained by mass action effects, where a reduction in enzymatic activity can be overcome by increasing the total amount of enzyme in the cell.

**Figure 2.**
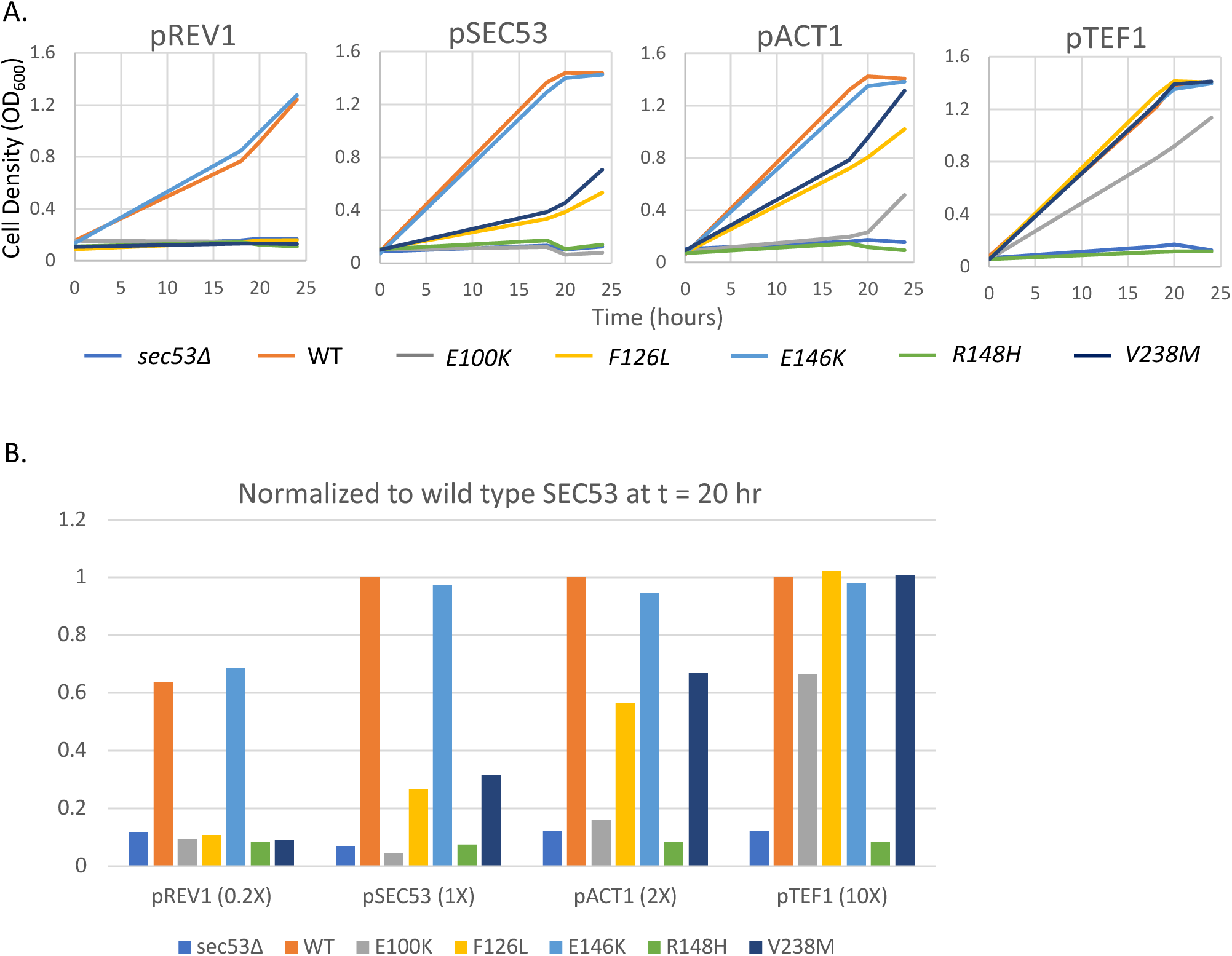
Comparison of growth of yeast SEC53 haploid alleles. (A) Graphs show growth of 10^-2^ dilution of OD 1.0 cells over time at 30°C in 50 µL SC+FOA media in a 384-well plate. 1X indicates the native SEC53 promoter. 2X indicates double the native promoter strength. 10X indicates 10X the native promoter strength. 0.2X indicates 20% of the native promoter strength. (B) Comparison of growth relative to wild type SEC53 at a single time point (t=18).

In contrast, reducing wildtype SEC53 mRNA expression to 20% of its native level with the REV1 promoter modestly, but consistently, compromises cell growth to 63.6% of the control (**Figure 2A and B**). This result shows that yeast cells are highly sensitive to the total amount of SEC53 protein. Interestingly, there was no toxicity associated with 100-fold over-expression of wildtype SEC53. Similar to wildtype, the E146K variant is defective only at the lowly expressing REV1 promoter (68.8%). This result suggests that the splicing defect of the 415G>A mutation in humans may reduce the abundance of still functional PMM2 E139K proteins to a sub-critical amount below the threshold for viability.

Comparing the strains relative to one another at a single time-point, we can see that the severity of growth defects of the SEC53 alleles correlates with the level of residual enzymatic activity of the PMM2 alleles reported using bacterial recombinant protein and PMM2-CDG patient fibroblasts (**Figure 2B**).

### SEC53 diploid variants recapitulate the growth of haploid variants

Most PMM2-CDG patients are compound heterozygotes with a recurring or novel mutation (usually missense) paired with the R141H null allele. The most common genotype is PMM2^F119L/R141H^. To further study genotype-phenotype relationships of the SEC53 variants, we generated homozygous and compound heterozygous diploids (**Figure 3**). As expected, a single copy of wildtype SEC53 is sufficient for normal diploid cell growth and there is no growth difference between homozygous wildtype and heterozygous SEC53^+/R148H^ (99.7%) cells (**Figure 3A**).

**Figure 3.**
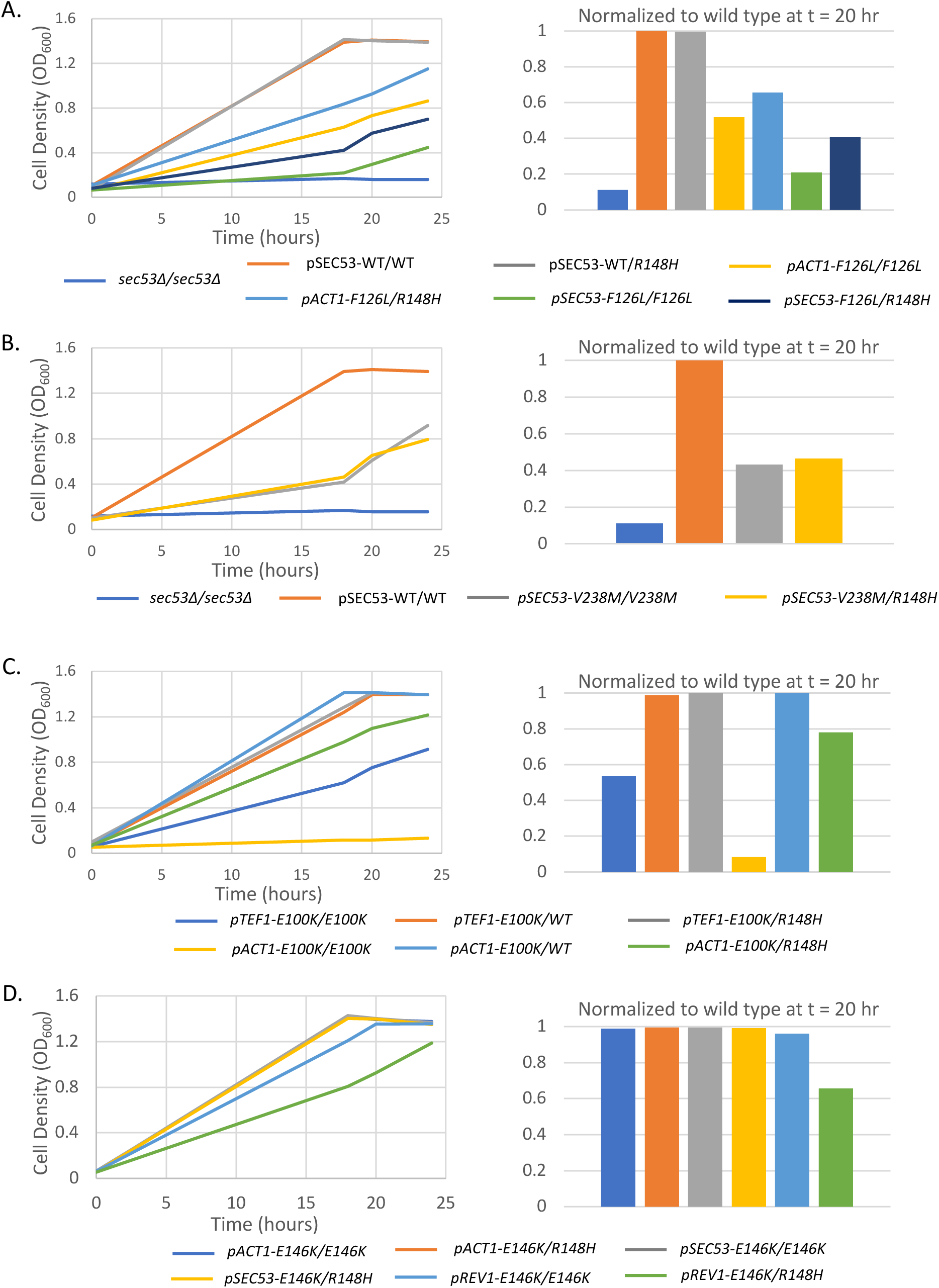
Comparison of growth of yeast SEC53 diploid alleles. Graphs show growth of 10^-2^ dilution of OD 1.0 cells over time at 30°C in 50 µL SC+FOA media in a 384-well plate. 1X indicates the native SEC53 promoter. 2X indicates double the native promoter strength. 10X indicates 10X the native promoter strength. 0.2X indicates 20% of the native promoter strength. Bar graphs show comparison of growth relative to homozygous wild type SEC53 diploids at a single time point (t=20). Strains with the indicated promoter are listed in the legend: A) SEC53-F126L, B) SEC53-V238M, C) SEC53-E100K, and D) SEC53-E146K.

Homozygous F126L diploids grow slower than their respective F126L/R148H heterozygous diploids (**Figure 3A**). In both 1X and 2X expression regimes, R148H heterozygosity improves growth. Normalizing to wildtype at 20 hours, the growth of pACT-F126L/F126L is 51.8% compared to 65.7% in pACT1-F126L/R148H. pSEC53-F126L/F126L is 20.9% compared to 40.7% in pSEC53-F126L/R148H. These results could be explained by the diminished ability of the PMM2 F126L variant to homodimerize, and the presence of R148H monomers allows formation of hemi-functional F126L:R148H heterodimers (Andreotti *et al.*, 2015). On the other hand, V238M causes protein misfolding but properly folded V238M monomers are competent to dimerize. The pSEC53-V238M/V238M homozygous diploid (43.2%) grows similarly to the pSEC53-V238M/R148H heterozygous diploid (46.4%) (**Figure 3B**). However, at the final 24 hour timepoint the pSEC53-V238M/V238M homozygous diploid was growing better than the pSEC53-V238M/R148H heterozygous diploid. This could be explained by the excess of R148H monomers relative to V238M monomers resulting in formation of nonfunctional R148H:R148H homodimers at a higher rate than hemi-functional V238M:R148H heterodimers.

Just as in the case of the dimerization-defective F126L variant, E100K homozygous diploids grew more poorly than E100K/R148H heterozygous diploids, which in turn grew more slowly than E100K/WT diploids (**Figure 3C**). Under the TEF1 promoter, E100K/E100K is 53.4% compared to 98.8% in E100K/WT and 100% in E100K/R148H. Under the ACT1 promoter, E100K/E100K is 8.2% compared to 100% in E100K/WT and 77.9% in E100K/R148H. Like F126, E100 is expected to affect dimerization and R148H may facilitate the formation of partially functional heterodimers. E146K, which is indistinguishable from wildtype, grew as expected except with the REV1 promoter: pREV1-E146K/E146K homozygous diploids (96.2%) grow better compared to the E146K/R148H heterozygous diploids (65.6%) (**Figure 3D**). The presence of R148H monomers results in fewer E146K:E146K homodimers which is still compatible with viability but not maximal growth rate.

### The phenotypes of human PMM2 alleles in yeast parallel SEC53 alleles

It was previously demonstrated that expression of human PMM2 rescues the lethality of a temperature-sensitive allele of SEC53, *sec53-6* (Hansen *et al*., 1997). We initially expressed human PMM2 cDNA but this failed to rescue yeast *sec53Δ*. For the first time we show that expression of human PMM2 cDNA that is codon-optimized for expression in yeast does in fact rescue *sec53Δ* (**Figure 4**). This suggests that the non-codon-optimized PMM2 was likely poorly expressed in yeast such that it was not sufficient for growth under the native SEC53 promoter. Subsequently, we expressed each of the human PMM2 variants in *sec53Δ* cells to determine whether PMM2 alleles behave the same as SEC53 alleles. Under the SEC53 promoter, PMM2 partially rescues *sec53Δ* to 71% of wildtype yeast SEC53 (**Figure 4A**). The degree to which each PMM2 variant rescues *sec53Δ* correlates with their reported residual enzymatic activity and supports the conclusion that the biochemical defect of each allele is conserved between yeast and humans. PMM2 E139K (68%) grows similarly to wildtype PMM2, in agreement with yeast SEC53 E146K. V231M compromises growth (55.6%) and F119L further compromises growth (44.8%). E93A, on the other hand, is not sufficient for growth under the SEC53 promoter and does not restore growth above *sec53Δ* cells. This is consistent with pSEC53-E100K and further supports the essentiality of E93 in PMM2 function.

**Figure 4.**
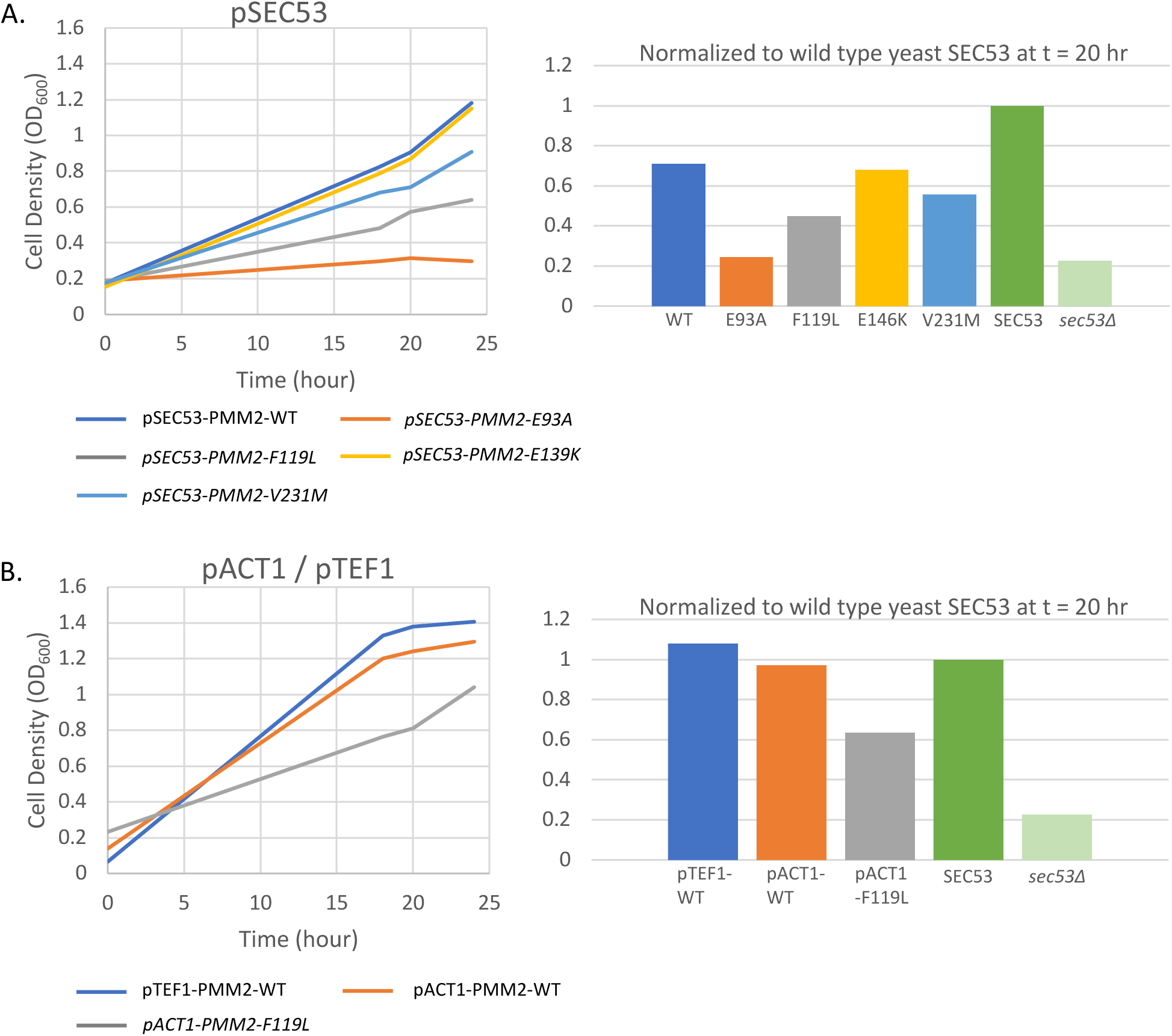
Expression of human PMM2 rescues growth of yeast *sec53Δ* cells. (A) Expression of the indicated human PMM2 variant under the SEC53 promoter. (B) Expression of the indicated human PMM2 variant under the ACT1 or TEF1 promoter. Left panels show growth over time and right panels show growth relative to wild type SEC53 at t = 20 hr.

Expressing PMM2 under the ACT1 promoter improves growth to 97.3% (**Figure 4B**). Expectedly, doubling the expression of F119L with the ACT1 promoter also improves its growth to 63.6%. Under the TEF1 promoter, PMM2 completely rescues *sec53Δ* cells. This suggests that PMM2 expressed in yeast may not be functioning optimally and requires higher expression levels. As is the case with SEC53, 10-fold over-expression of PMM2 is not cytotoxic. Based on these results, we can characterize the deficiency of a given PMM2 allele in yeast based on its effect on cell growth over a range of protein expression regimes and in different compound heterozygous combinations.

### Measuring phosphomannomutase activity levels in SEC53 variants

We performed a PMM2 enzymatic assay to determine whether the expression level of the variants correlates with the enzymatic activity measured in wildtype and mutant yeast cell lysates. We assayed phosphomannomutase (PMM) activity by measuring the reduction of NADP+ to NADPH spectrophotometrically at 340 nm with a plate reader (Van Schaftingen & Jaeken, 1995; Pirard *et al.*, 1999; Kjaergaard *et al.*, 1999; Sharma *et al*. 2011; and Citro *et al.*, 2017). NADPH production is achieved by a series of four enzyme-coupled reactions shown in **Supplemental Figure 1**. Mannose-1-phosphate and NADP+ are provided as substrates for the reactions along with phosphomannose isomerase, phosphoglucose isomerase, glucose-6-phosphoate dehydrogenase, and PMM2 from the protein lysate. Additionally, glucose 1,6-bisphosphate is used as an activator of PMM2 activity.

We determined that wildtype SEC53 expression level and growth correlates to the PMM activity detected by the assay (**Figure 5A and B**). Lysates from pTEF1 and pACT1 cells consistently show elevated activity compared to pSEC53 cells. Lysates from pREV1-SEC53 cells show less activity (70%), consistent with its growth relative (64%) to pSEC53-SEC53. For V238M, the expression level correlates with enzymatic assay and with growth, but not to the extent expected (**Figure 5C**). For example, pTEF1-V238M rescues growth completely (100%), but PMM activity in these cells are below half of wildtype cells (37.5%) (**Figure 5C**). pACT1-V238M also rescue growth to a higher degree than the enzymatic activity (29%) of these cells would indicate. This is also true for F126L (**Figure 5C**). However, pACT1-F126L (19.4%) and pSEC53-F126L (17.4%) does not show significant difference in PMM activity. This indicates that for SEC53 mutations, either mannose-1-phosphate levels or PMM enzymatic activity in cell lysates does not necessarily reflect the growth of these cells. In other words, SEC53 variants may only require a modest boost in PMM enzymatic activity and/or mannose-1-phosphate levels above their baselines to achieve maximal growth rate and presumably a fully glycosylated proteome.

**Figure 5.**
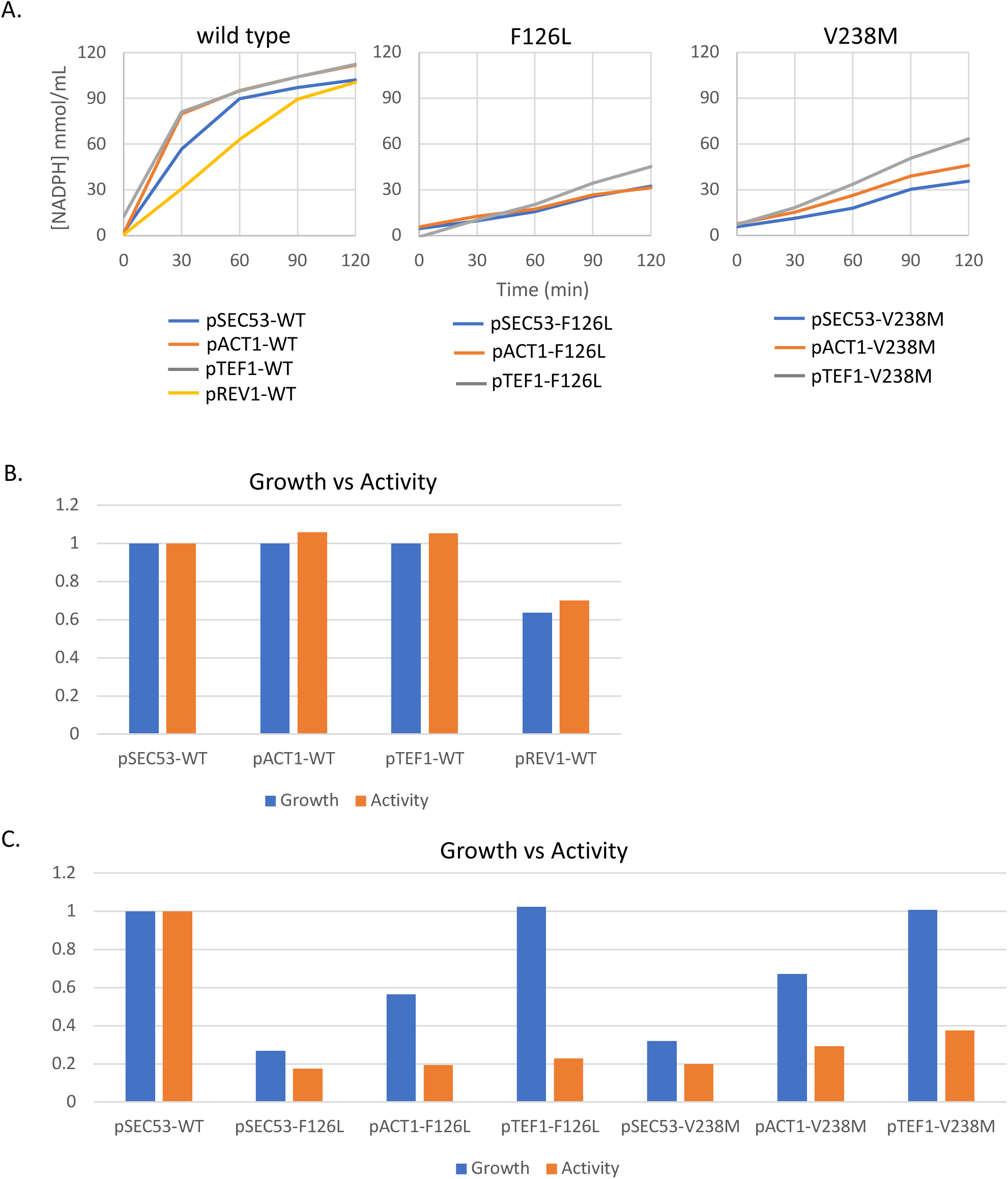
Phosphomannomutase activity of SEC53 alleles. (A) Graphs show NADPH formation over time in 200 µL reaction with 0.1 µg/mL protein lysate from the strains indicated. (B) Phosphomannomutase activity of the indicated SEC53 variant relative to pSEC53-WT plotted against cell growth relative to pSEC53-WT.

### Drug repurposing screen in yeast models of PMM2-CDG identified three novel chemical modifiers

We advanced pACT1-F126L, pSEC53-V238M, and pSEC53-F126L haploids and pACT1-F126L/pACT1-R148H, pSEC53-V238M/pSEC53-R148H, and pSEC53-F126L/pSEC53-R148H heterozygous diploids to high-throughput drug screens. Screening was done with a 2,560 compound MicroSource Spectrum library consisting of FDA approved drugs, bioactive tool compounds, and natural products. Each strain was screened in duplicate in 384-well plates, with each plate containing 32 wells of the negative control (no drugs) and 24 wells of the positive controls (wildtype cells). As shown in **Figure 6A**, the positive and negative controls showed good separation of Z-scores, which allowed us to distinguish even a relatively modest rescue of growth in the screen. Additionally, there is good correlation between replicates (correlation >0.4 for all panels) (**Figure 6B**). With a Z-score cutoff of 2.0, we identified six “pre-hit” compounds that looked promising because their ability to rescue growth is conserved between the haploid and diploid strains (**Figure 6C**).

**Figure 6.**
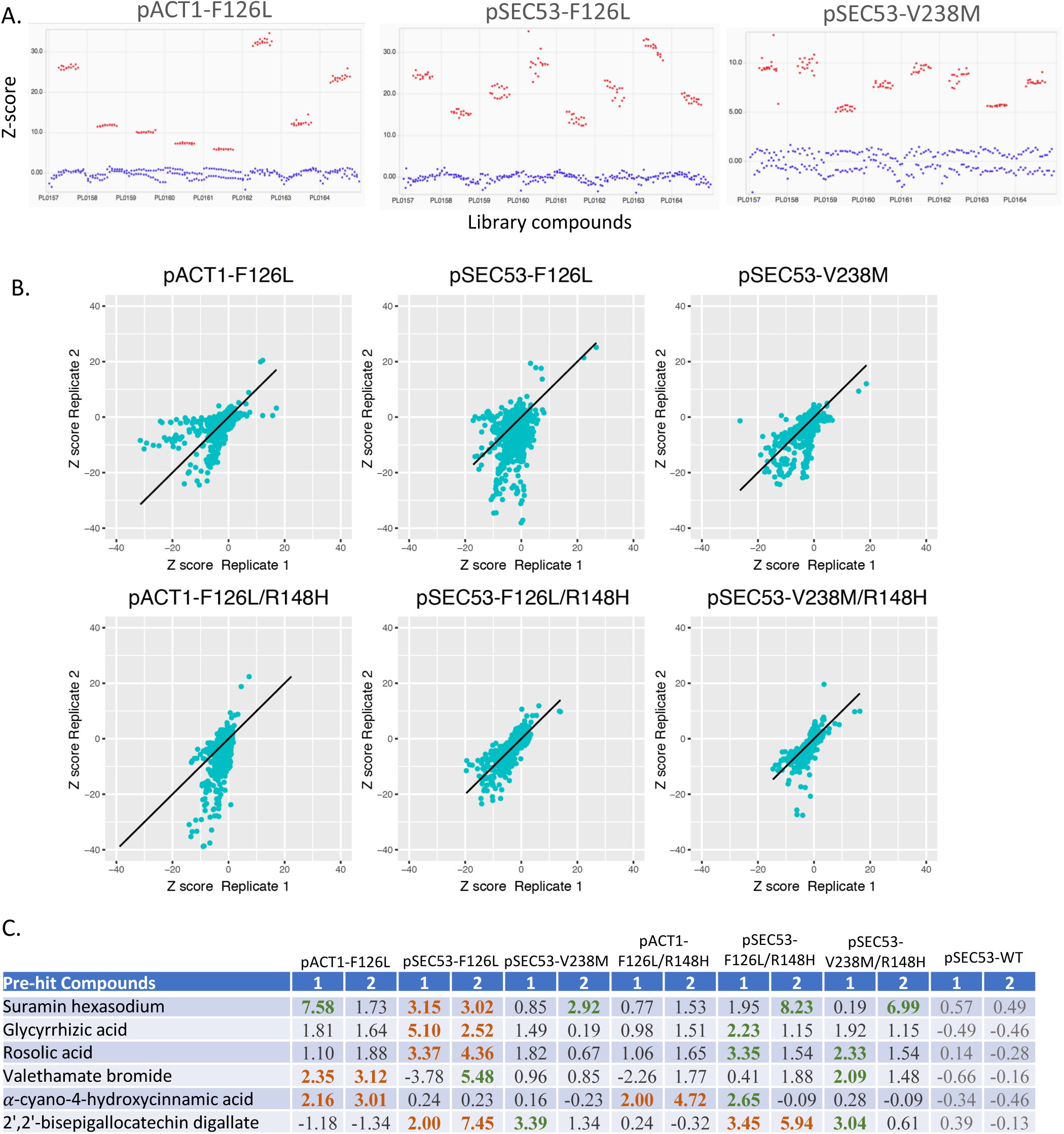
Summary of the 2,560 drug repurposing screen. (A) Comparison of z-scores between the negative vs positive controls of representative datasets show separation of data. (B) Comparison of z-scores between replicates show positive correlation between duplicate datasets. (C) Pre-hit compounds and z-scores. We identified six compounds from the Microsource Spectrum library that showed a z-score of ≧2.0 in growth in at least 2 replicates of the same allele (orange). Green indicates a z-score of ≧2.0, but did not replicate in the duplicate screen.

We reordered these compounds from the vendor as dry powder to retest them in multi-point dose response assays. We found that three of the six compounds showed consistent and dose-dependent rescues in haploid and heterozygous diploid cells (**Figure 7**). The negative control compound cysteamine hydrochloride did not rescue at any dose (**Figure 7D**). At the maximum dose tested, alpha-cyano-4-hydroxycinnamic acid rescues the F126L allele by 17.7% in haploid and 24.5% in diploid cells. Interestingly, this rescue is specific to the 2X pACT1 expression level (**Figure 7A** and **Supplemental Figure 2**).

**Figure 7.**
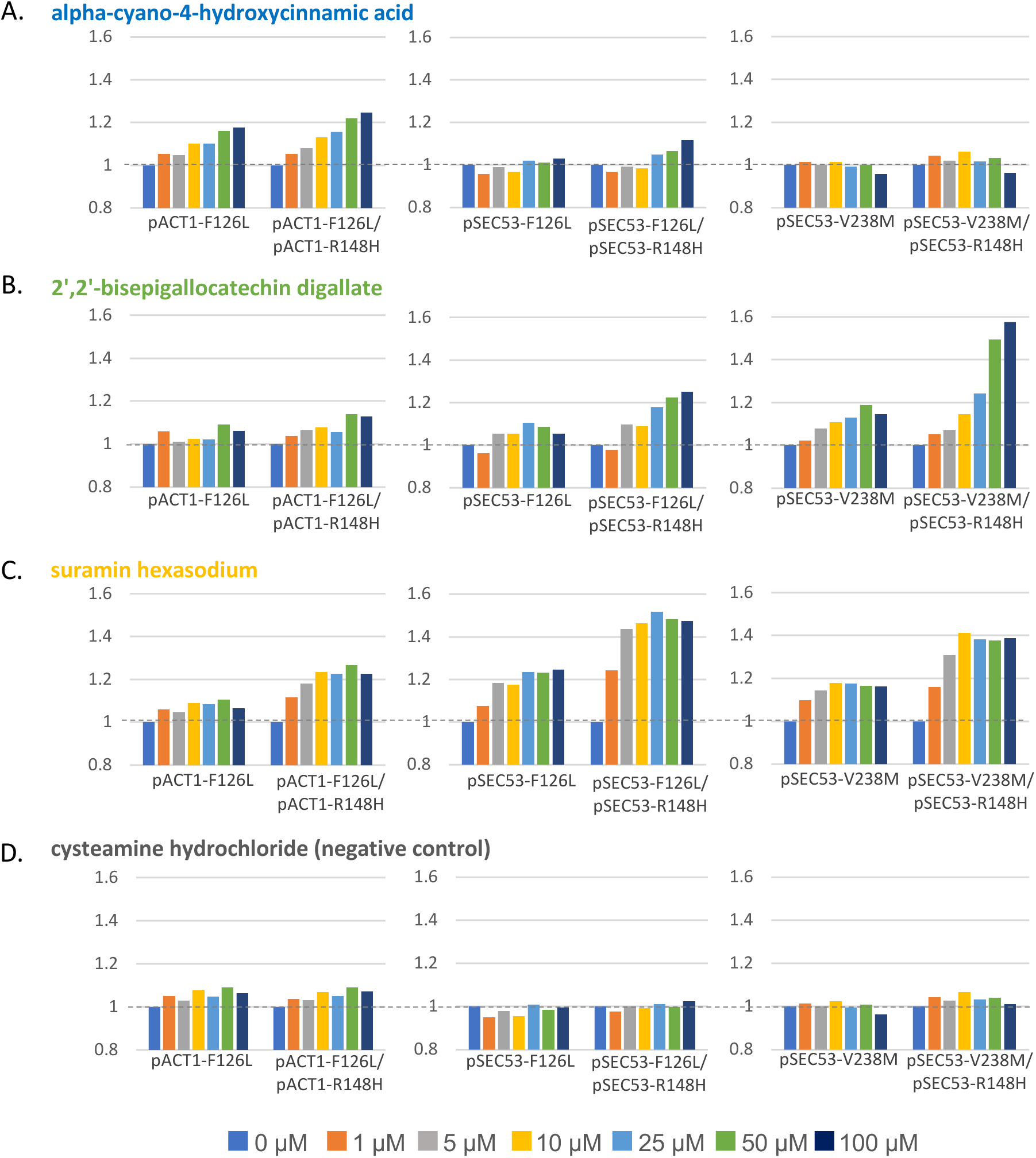
Three compounds show differential rescue of growth of SEC53 variants. (A) Growth of cells in alpha-cyano-4-hydroxycinnamic acid at the indicated dose relative to growth in the absence of compound (0 µM). (B) Growth of cells in 2’,2’-bisepigallocatechin digallate at the indicated dose relative to growth in the absence of compound (0 µM). (C) Growth of cells in suramin hexasodium at the indicated dose relative to growth in the absence of compound (0 µM). (D) Growth of cells in cysteamine hydrochloride (negative control) at the indicated dose relative to growth in the absence of compound (0 µM).

2’-2’-bisepigallocatechin digallate rescues growth of pSEC53-F126L and pSEC53-V238M cells, and pACT1-F126L to a much lesser extent (**Figure 7B**). It also exerts a stronger response in heterozygous diploid cells than haploids. Growth of pSEC53-F126L improved by as much as 10.6% and pSEC53-F126L/R148H by 25%. pSEC53-V238M improved by 14.6% and pSEC53-V238M/R148H by 57.7%. pACT1-F126L cells only improved by 6.3% and 12.9% in pACT1-F126L/R148H (**Figure 7B**).

Similarly, suramin hexasodium also rescues growth of the pSEC53 variants more than the pACT1 variants, and diploids more strongly than haploids (**Figure 7C**). Growth of pSEC53-F126L improved by 24.5%, pSEC53-F126L/R148H by 47.4%, pSEC53-V238M by 16.1%, and pSEC53-V238M/R148H by 38.9%. In contrast, pACT1-F126L cells showed 6.7% improvement and pACT1-F126L/R148H showed 22.6%.

In summary, we found that alpha-cyano-4-hydroxycinnamic acid specifically rescues pACT1-F126L cells, while 2’-2’-bisepigallocatechin digallate and suramin hexasodium preferentially rescue pSEC53-F126L/R148H and pSEC53-V238M/R148H heterozygous diploid cells.

### Discussion

We created new yeast models of the congenital disorder of glycosylation PMM2-CDG and demonstrated their relevance toward developing therapeutics in four ways. First, yeast-codon-optimized human PMM2 fully rescues the lethality of *sec53Δ* cells. Second, each of the five pathogenic PMM2 missense mutations studied herein and their orthologous SEC53 mutations support comparable yeast cell growth in a *sec53Δ* whole-gene deletion background. The rank-ordered severity of the mutation pairs from weakest to strongest loss-of-function is: E139K & E146K >> V231M & V238M > F119L & F126L >> E93A & E100K >> R141H & R148H. Third, the residual enzymatic activities reported in the literature for the five PMM2 variants are correlated to the extent of yeast cell growth rescued by these variants. Fourth, a pilot drug repurposing screen successfully identified three compounds – an FDA approved drug, a tool compound and a GRAS (generally recognized as safe) plant-based natural product – that appear to act by two different mechanisms of action.

The three most well studied variants – R141H, F119L and V231M – all behaved as expected in haploid and homozygous diploid cells. A residual PMM2 enzymatic activity is not reported for bacterially expressed recombinant E93A protein, and our results clearly indicate the strong loss-of-function of both E93A and E100K variants in haploid and homozygous diploid cells. Contrary to the reported 25% residual PMM2 enzymatic activity of the E139K variant, our results indicate that both E139K and E146K variants grow similarly to wildtype controls with only a modest growth defect evident at the lowest expressing REV1 promoter, or 20% of the native SEC53 promoter strength. The two dimerization-defective variants F126L and E100K grew better when in trans with R148H than as homozygotes because R148H monomers can dimerize with either F126L or E100K monomers to form hemi-functional heterodimers. On the other hand, the known strongly folding-defective variant V238M and the weakly hypomorphic variant E146K grew better in homozygosity than in trans with R148H. We speculate that a larger pool of R148H monomers relative to V238M or E146K monomers due to the greater stability of the latter results in a higher proportion of nonfunctional R148H:R148H homodimers diluting the total pool of fully functional V238M or E146K homodimers. Collectively, these results suggest two different therapeutic approaches, one for when a PMM2 allele is dimerization defective and another for when a PMM2 allele is folding defective or prone to aggregation and rapid turnover.

Based on the results of the phosphomannomutase enzymatic assay we adapted for yeast cell lysates, we challenge the reported residual enzymatic activities of F119L and V231M as 25% and 37.5% of control levels, which were determined using bacterially expressed recombinant PMM2 variants. Using wildtype SEC53 expressed at 20% of endogenous protein levels as a benchmark, we would predict that the residual phosphomannomutase activity of V238M is ∼10% of control levels and that of F119L is ∼5% of control levels. In our hands, PMM2^F^119^L/R141H^ patient fibroblast cells show 10% enzymatic activity. That would imply that the residual enzymatic activity of the E93A variant may be less than 5% of control levels, while the residual enzymatic activity of the E139K variant may be greater than 25% of control levels.

The pharmacology of the three hits from the Microsource Spectrum screen suggests possible explanations for rescue of PMM2-CDG involving antioxidant effects and perturbation of glycolysis. alpha-cyano-4-hydroxycinnamic acid has been described as a monocarboxylate transporter inhibitor (Diers *et al.*, 2012) as well as an aldose reductase inhibitor (Zhang *et al*., 2016). 2’-2’-bisepigallocatechin digallate is also known as theasinensin, an active compound in oolong tea, and has numerous antioxidant properties shared by other plant-based polyphenols (Butt *et al.*, 2014). Suramin is an FDA approved drug for African sleeping sickness and river blindness, and its mechanism of action against trypanosomes involves perturbations to N-acetylglucosamine biosynthesis (Alsford *et al.*, 2012). Future experiments include testing whether any of these three compounds rescue PMM2 enzymatic activity in patient fibroblasts.

These yeast based PMM2-CDG patient avatars not only can be used to identify potential therapeutics demonstrated in this study, but can also be used in genetic modifier screens, genetic interaction screens, and basic studies to elucidate the pathophysiological cascade of PMM2-CDG starting at the cellular level and working up in model organism complexity.

## Acknowledgements

We thank Maggie’s PMM2-CDG Cure, LLC and the Carmichael family for funding this work. We thank Next Interactions for the initial construct engineering and growth experiments, and Dr. Sangeetha Iyer for comments and feedback on the manuscript.

---

**Supplemental Figure 1.**
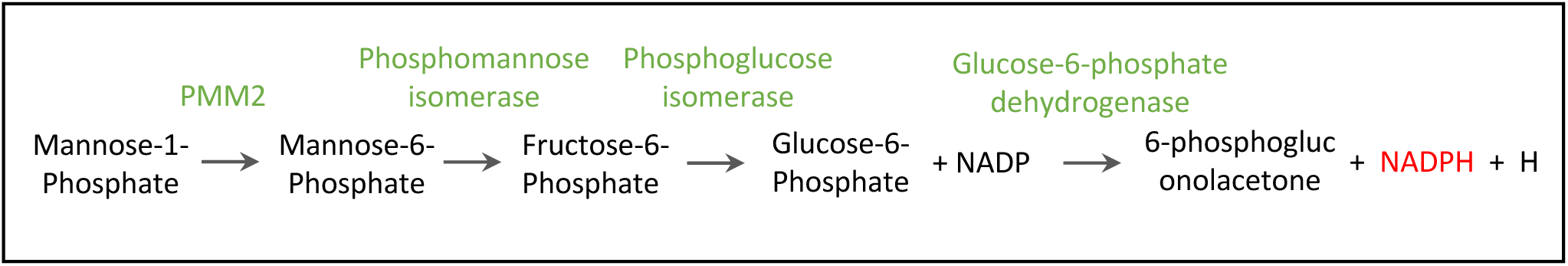
Phosphomannomutase enzymatic assay reactions.

**Supplemental Figure 2.**
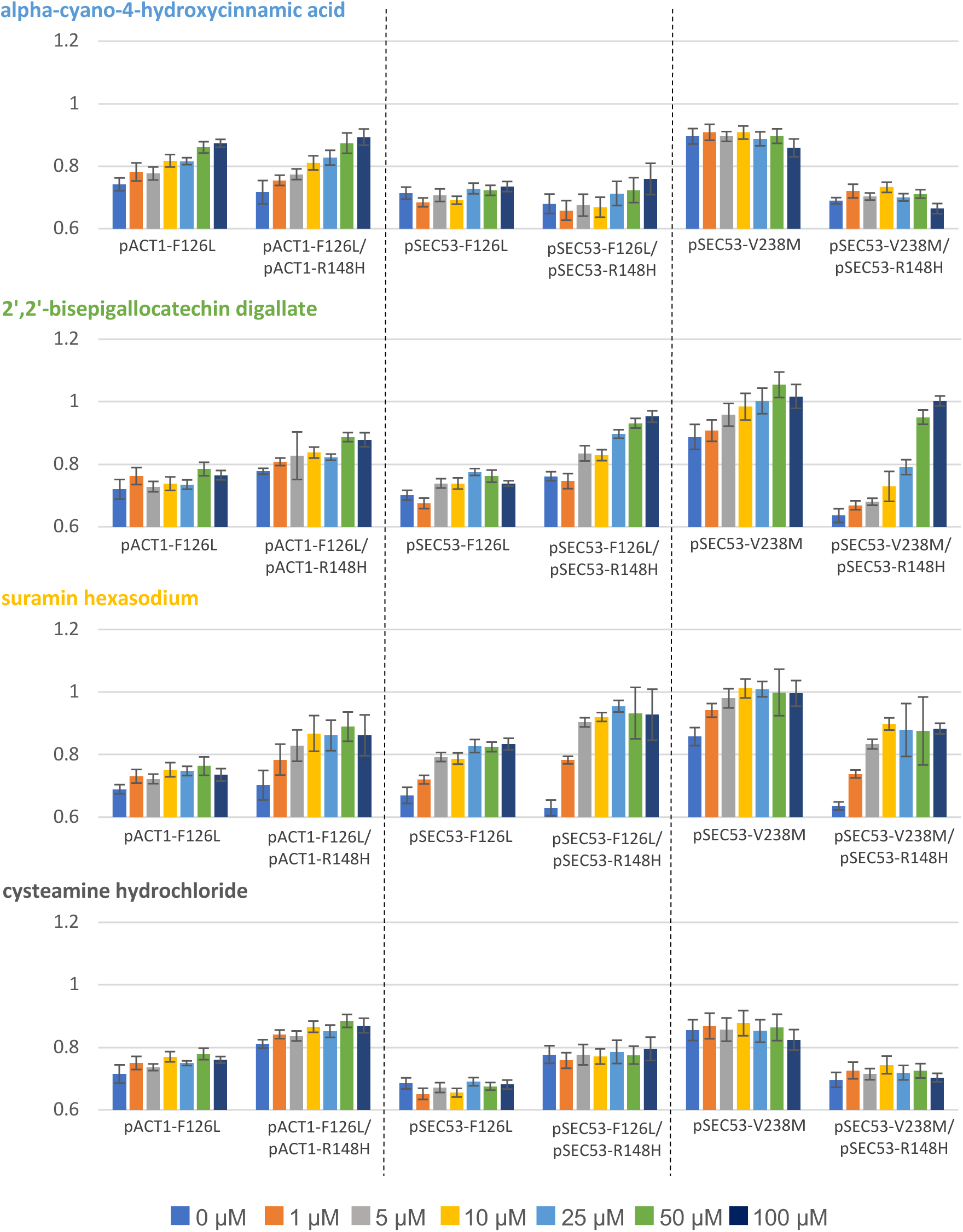
Data shows the mean of the cell density (OD_600_) across 16 samples of the each SEC53 variants grown in the indicated compound and dose.

**Supplemental Table 1.**
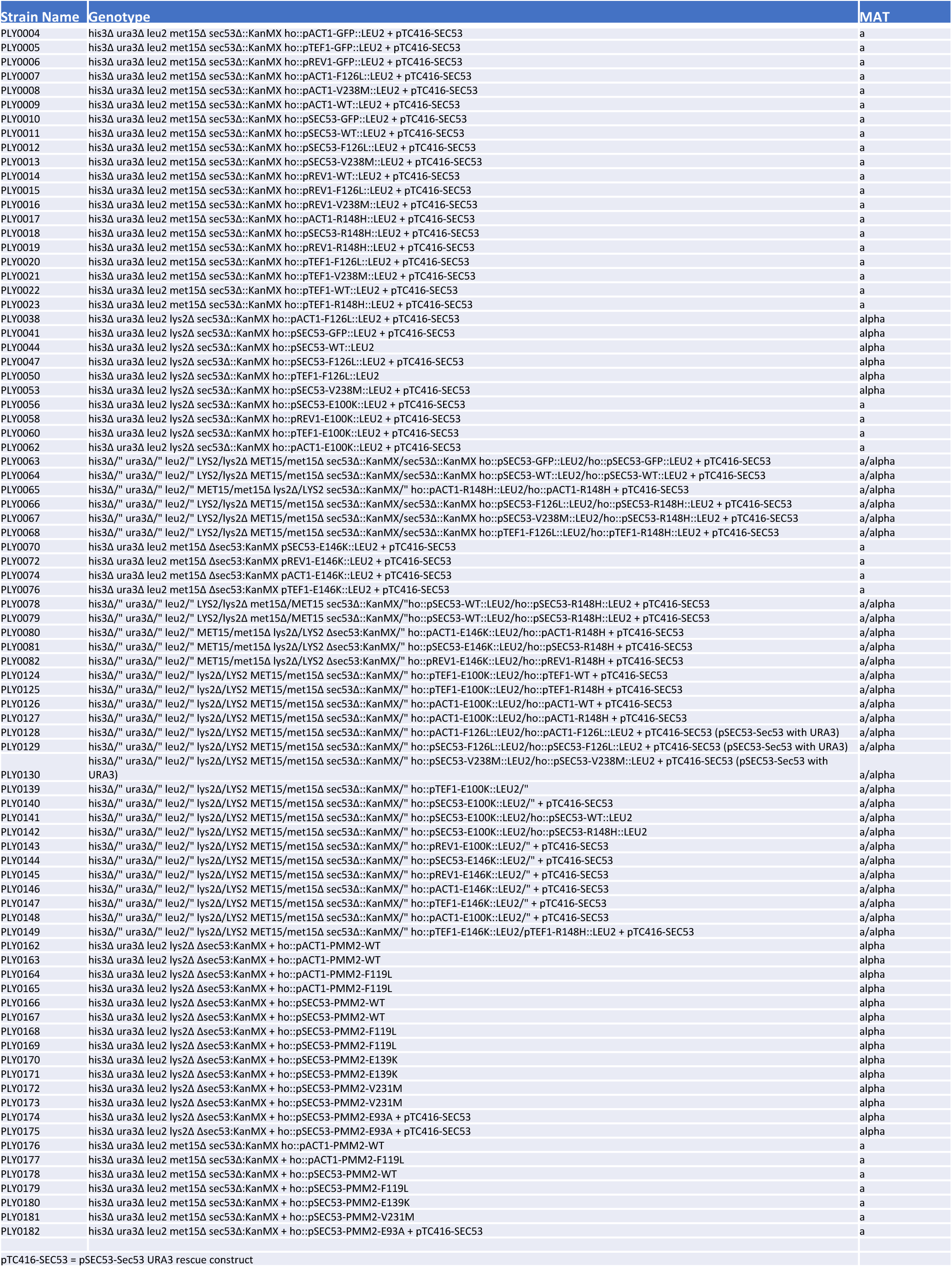
List of strains used in this study. Strains are available upon request.

